# Host genotype shapes root mycobiota in durum wheat

**DOI:** 10.1101/2024.10.27.616629

**Authors:** Margot Trinquier, Michel Colombo, Hélène Fréville, Jacques David, Aline Rocher, Benoit Lefebvre, Christophe Roux

## Abstract

- In addition to environmental factors, plant genetics play a key role in shaping the root microbiota; however, the extent of this genetic control remains underexplored. Using a collection of 181 wheat lines derived from a genetically diverse population, we investigated the influence of wheat genotypes on the composition of the root endophytic mycobiota and explored the genetic determinants driving these relationships.
- We first characterized the mycobiota associated with the roots of field-grown lines from the Evolutionary Prebreeding pOpulation (EPO) using Internal Transcribed Spacer 2 (ITS2) barcoding. Fungal diversity was then correlated with wheat genetics by quantitative methods, including heritability analysis and Genome Wide Association Studies (GWAS), to identify novel genetic determinants influencing the mycobiota composition.
- Fungal species richness showed a positive correlation across most fungal clades, except between *Mortierellomycotina* and *Glomeromycotina*. Some specific fungal clades, such as *Olpidiomycota* or *Chytridiomycota*, underscored their potential as root endophytes. Additionally, we observed higher heritability in fungal clades (*i.e.* at the phylum or subphylum rank) that exhibit a homogenous trophic mode, such as the biotrophic Arbuscular Mycorrhizal Fungi (AMF).
- This study identifies 11 QTLs associated with mycobiota composition at the clade level. By shedding light on the genetic control of fungal diversity and uncovering key fungal associations, this work enhances our understanding of plant-microbiota interactions and highlights the potential for breeding strategies to optimize these relationships.

## Introduction

Plants establish complex interaction networks with microorganisms that play a crucial role in enhancing their nutrition and resistance to biotic and abiotic stresses (Trivedi *et al*., 2020). When encountering soil microbiota, root systems orchestrate their microbial environment, spanning from the rhizosphere and rhizoplane to the root endosphere (van der Heijden & Schlaeppi, 2015). The relative importance of plant genetics and soil environmental determinants on the formation of root microbial communities is still the subject of debate. Numerous abiotic factors have been shown to create local ecological niches that impact soil microbiome. In agricultural settings, these factors encompass soil physico-chemical properties, water and nutrient availability, climatic conditions, agricultural practices, and cropping history (Lupwayi *et al*., 1998; Hartman *et al*., 2018; Azarbad *et al*., 2020). Beyond these abiotic factors, the genetics of the host plant also exerts influence on the rhizosphere, rhizoplane and root endosphere microbiomes (Zhang *et al*., 2023).

Plant genetics has been identified as a key driver of phyllosphere microbiome variation (Horton *et al*., 2014; Wallace *et al*., 2018; Beilsmith *et al*., 2019; Roman-Reyna *et al*., 2020). However, assessing such correlation in roots is inherently more complex. First, the high diversity of soil microbes challenges the robustness of diversity analyses. Secondly, roots harbor multiple associated microbiomes (rhizosphere-rhizoplane-endosphere), influenced by various intertwined factors such as root exudate composition (involving host-microbe chemical signaling and nutrient exchange) and plant immune system (Zhang *et al*., 2023). Nevertheless, studies conducted on various plant species, including crops, grown under either controlled or field conditions, with or without abiotic stresses, have successfully identified correlations between plant genetics and root microbiomes (Schlaeppi *et al*., 2014; Edwards *et al*., 2015; Deng *et al*., 2021; Escudero-Martinez *et al*., 2022; Wang *et al.,* 2022; Andreo-Jimenez *et al.,* 2023; Roux *et al*., 2023).

Among the different phyla of organisms that make up the microbiome, much more is known about bacteria than fungi. However, the plant fungal microbiota, or mycobiota, plays a critical role in plant fitness (Bonfante *et al*., 2019). The Fungal kingdom comprises approximatively 3.8 million species (Hawksworth & Lücking, 2017), categorized into several phyla The most diverse of these are *Ascomycota*, *Basidiomycota* and *Mucoromycota*, the latter of which includes three subphyla: *Mucoromycotina*, *Mortierellomycotina* and *Glomeromycotina* (Spatafora *et al*., 2016; Li *et al*., 2021). Soil represents the largest reservoir of fungal diversity (Tedersoo *et al*., 2021). This diversity decreases near plant roots and further within the roots as endophytic fungi (Urbina *et al*., 2018), as a result of successive selective processes (Mesny *et al*., 2021). Endophyte fungi were first identified as ascomycetes (Rodriguez *et al*., 2009), but barcoding investigations based on high-throughput sequencing have greatly enlarged our vision on their diversity (Harrison & Griffin, 2020). Indeed, recent findings indicate that many taxa are currently found as part of the root mycobiota and are even proposed as plant growth promoting fungi (Zhang *et al*., 2020). Interestingly, many fungi previously classified as strict saprotrophs have been identified as root endophytes through barcoding, suggesting a need to redefine their ecological roles (Põlme *et al*., 2020).

In wheat, one of the world’s major crops, as in other plant species, the root mycobiota has received much less attention than its bacterial counterpart (Donn *et al*., 2015; Naylor *et al*., 2017; Rossmann *et al*., 2020; Simonin *et al*., 2020; Kavamura *et al*., 2021). Nevertheless, using 12 cultivars grown in pots, Latz and co-authors found that bread wheat genetics significantly affects the root mycobiota (Latz *et al*., 2021). This underscores the importance of identifying genomic regions that govern the composition of the wheat root mycobiota.

Durum wheat was first domesticated 10,000 years ago in the Fertile Crescent and then bred and selected for agronomic characteristics. As a result of recurrent selective events during domestication and breeding, genetic diversity of wheat and more broadly of grass crops has been strongly reduced (Glémin & Bataillon, 2009; Maccaferri *et al*., 2019). Such loss might have altered the ability of wheat crops to interact with beneficial microorganisms (Sawers *et al*., 2008; Fréville *et al*., 2022). In this context, studying the genetic determinants underlying root mycobiota organization could help designing selection schemes for improving beneficial microbial interactions.

Deciphering the genetic architecture of traits has been tremendously accelerated by the use of Genome Wide Association Studies (GWAS), including multivariate analysis as traits (Zhang *et al*., 2018). The root mycobiota is currently investigated by metabarcoding through amplification and sequencing of specific genomic regions (Tedersoo *et al*., 2022). Integrating these two approaches would enable the identification of the host genetic determinants that underpin the organisation of the root mycobiota. (Zancarini *et al*., 2021). However, as metabarcoding methods allow detecting hundreds of taxa in a sample, comparison of mycobiota composition between samples has thus to be addressed with summarized metrics as diversity indexes and multivariate analysis used to describe the samples in a much more simplified numerical space. Diversity indices and genotype coordinates following multivariate analysis can thus be considered as quantitative traits on which the relationship with host genetics can be investigated by GWAS.

To avoid establishing spurious genetic associations, the study of mycobiota on a large diversity of host genotypes in field requires experimental designs with replicates and adequate spatial modeling to control intra-field spatial variability due to micro-environment. Such models have been recently developed in agronomy (Rodríguez-Álvarez *et al*., 2018), and have been successfully applied in a field-based study of sorghum rhizosphere microbiota (Deng *et al*., 2021). The partially replicated design (p-rep design) is particularly adapted to these issues as it enables to find a trade-off between estimation of environmental variation and experimental efforts (Smith *et al*., 2006). Yet, this type of experimental setup has not yet been used to investigate the genetic determinants of root mycobiota composition.

In this study, we performed fungal metabarcoding approaches and GWAS on a collection of 181 field-grown durum wheat inbred lines from the highly diversified Evolutionary Prebreeding durum wheat pOpulation (EPO - David *et al*., 2014). As the genetic diversity in durum wheat has been strongly reduced since the domestication of the wild relatives, wild and domesticated emmer wheat lines were crossed in this dynamic population to enrich in allelic diversity. We here address two key questions: (1) Is the ability of durum wheat roots to associate with fungi genetically determined? (2) Which fungal phyla are most strongly influenced by wheat genetic control?

## Material and Methods

### Genetic material

The Evolutionary Prebreeding durum wheat pOpulation (EPO) was obtained by intercrossing a durum wheat composite cross segregating for a spontaneous male sterility with some from tetraploid taxa, including wild, primitive, and domesticated emmer wheat. It has been developed since 1997 at INRAE Montpellier, France. After twelve generations of open pollination, a set of 181 EPO lines was extracted and fixed by successive generations of selfing (David *et al*., 2014). These lines were genotyped with the TaBW410K marker high-throughput genotyping array (Rimbert *et al*., 2018), from which marker and presence/absence variants (Off Target Variants) were extracted. This set of lines has a weak linkage disequilibrium (LD) structure, making this panel ideal for GWAS. Non-polymorphic markers and markers with a minor allele count under 10 were discarded. Missing values were imputed as the observed allele frequencies at each locus. After these steps, 46,439 markers were available for further analysis.

### Experimental design

We grew the 181 EPO inbred lines in a 0.10 ha field at Mauguio (43.6208218; 3.9851894, experimental unit DIASCOPE, INRAE) in the south of France during the 2020-2021 season. Each plot consisted of 3 rows, each 1.5 m long spaced 0.23 m apart (Fig. **1a,b**). Each inbred line was replicated twice in plots randomly distributed across two blocks (Fig. **1c**). Seeds were sown on October 19, 2020, with 70 seeds per row. Plots were managed using shallow tillage, mechanical and hand weeding. Organic fertilization (Orga VIO 10.5.0, Violleau at 600 kg per ha) and irrigation were supplied to prevent resource limitations.

**Figure 1:**
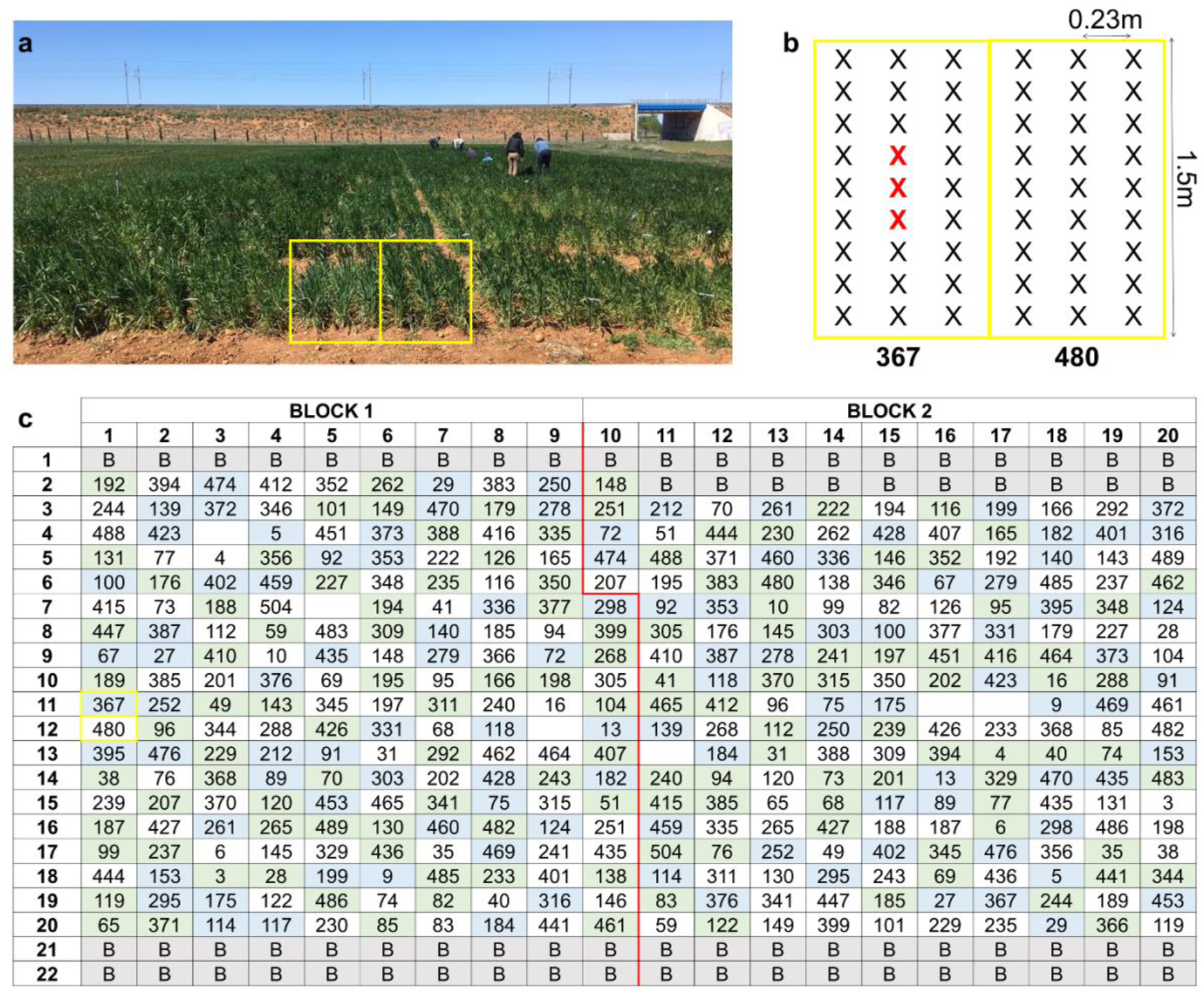
(**a**) Picture of the field trial taken at the time of the sampling. The plots containing EPO inbred lines number 367 and 480 are delimited by yellow rectangles. (**b**) Plot sampling design: a sample consists of a pool of root systems from at least three plants (shown in red) in the middle of the central row. (**c**) Trial design in two blocks delineated with the red line, arranged in 22 lines and 20 columns and surrounded by a border (B) of elite durum wheat. Inbred line identification number is shown for each plot. Plots colored in green were sampled once, whereas those in blue were sampled twice into each block. The two EPO inbred lines shown in (**a**) and (**b**) are boxed in yellow.

Root samples were collected in April 2021 before the heading stage. Each sample consisted of a pool of root systems up to approximately 15 cm depth and belonging to a minimum of three plants. Sampling was conducted at the center of the central row within each plot to minimize border effects (Fig. **1b**). For 128 out of the 181 lines, root samples were randomly collected between the two blocks. For the remaining 53 lines, root samples were collected in each block. This sampling design corresponds to a p-rep design, where 30% of the test entries were replicates, enabling the calculation of heritability values while considering spatial effects.

### Characterization of mycobiota community

#### DNA extraction

Samples were carefully hand washed with tap water to remove all aggregating soil, using 180 µm sieves to collect fine root fragments. Cleaned roots were frozen in liquid nitrogen and stored at −80 °C before DNA extraction with a ZymoBIOMICS 96 DNA Kit in plate (Zymo Research). Two grinding steps were performed: a first manual grinding of the entire sample with mortar and pestle, and a second on a fraction of root powder in tubes containing beads provided with the kit, in a mixer mill MM400 (Retsch). DNA extraction was performed following the supplier protocol. DNA samples were then stored at −20°C.

#### Amplicon library and sequencing

The ITS2 rRNA amplicon library was generated using a nested PCR. Two primer pairs were used: 1) pair rcAMDGR (5’-ATGATTAATAGGGATAGTTGGG-3’) (Sato *et al*., 2005) & ITS4 (5’-TCCTSCGCTTATTGATATGC-3’) (White *et al*., 1990) that does not amplify wheat ITS2 rRNA, and then, 2) pair ITS 86F (5’-GTGAATCATCGAATCTTTGAA-3’) (Turenne *et al*., 1999) & ITS4 that produces an amplicon size compatible with Illumina MiSeq. For both PCR, the GoTaq G2 DNA polymerase (Promega) was used in a mix with the following final concentration: for PCR 1, 1X GoTaq Reaction Buffer, 0.08mM dNTPs, 0.32µM of both primers, 0.04U/µl of polymerase and 2% DMSO, and for PCR 2, 1X GoTaq Reaction Buffer, 0.2mM dNTPs, 0.4µM of both primers, 0.02U/µl of polymerase and 2.5 mM of MgCl2. Nuclease-Free Water for Molecular Biology (Sigma-Aldrich) was used for each PCR mix. PCR was performed with an initial denaturation at 95°C for 5 min and final elongation at 72°C for 10 min; for PCR 1, 40 cycles were used at 95°C for 5 sec - 62°C for 30 sec - 72°C for 1 min; and for PCR 2, 35 cycles at 95°C for 30 sec - 55°C for 30 sec - 72°C for 1 min. Amplification of ∼300 bp was visualized thanks to electrophoresis on 2% agarose gel.

Libraries were generated using the Illumina two-step PCR protocol and normalized using SequalPrep plates (Thermofischer). Paired-end sequencing with a 2×250 bp read length was performed at the Bio-Environment platform (University of Perpignan Via Domitia Perpignan, France) on a MiSeq system (Illumina) using v2 chemistry according to the manufacturer’s protocol. Data were deposited at SRA (PRJNA1070864).

#### Read processing

Raw reads were processed with the pipeline FROGS version 3.2.3 (Bernard *et al*., 2021) using the Galaxy interface (https://vm-galaxy-prod.toulouse.inrae.fr/galaxy_main/). Reads were merged, pre-quality filtered, dereplicated and trimmed at 400 bp with Vsearch version 2.17.0, flash version 1.2.11 and cutadapt version 2.10. Then clustering was done using the SWARM algorithm (Mahé *et al*., 2014) with an aggregation distance of 1 allowing to build very fine clusters with minimal differences. Chimeras and singletons were removed. Filtering was done on a minimum proportion to keep a zero-radius Operational Taxonomic Unit (zOTU) with at least 0.005% of all sequences, i.e. represented by at least 261 reads for the whole sampling (Escudié *et al*., 2018). Then ITSx filter version 1.1.2 was used to keep only sequences that correspond to the ITS2 region. Taxonomy affiliation was performed by BLAST using the UNITE Fungi 8.3 database (Kõljalg *et al*., 2019; Nilsson *et al*., 2019; Urbina *et al*., 2019). Rarefaction curves were done for each sample in order to detect sequencing quality. From these data, we extracted presence / absence data of each zOTU within each sample. To describe mycobiota community composition, we performed 6 independent multiple correspondence analysis (MCA) on presence / absence of each zOTU using the *MCA* function of the FactomineR package (Lê *et al*., 2008), one for each of the 6 fungal clades identified by the metabarcoding approach (Zancarini *et al*., 2021). We plotted either zOTUs or genotypes and extracted each coordinate along the MCA axes.

We also computed three alpha diversity indices for each EPO line, the *Observed*, *Shannon* and *Inverse Simpson* indices, using the R package *phyloseq* (McMurdie & Holmes, 2013). These indices were calculated for each fungal clade, referring either to a fungal phylum or a fungal subphylum. *Observed index* corresponds to the zOTU richness, i.e. the number of observed zOTUs in a given sample (Spellerberg & Fedor, 2003). *Shannon index* represents both richness and relative abundance of one zOTU among all (Shannon & Weaver, 1949). *Inverse Simpson index* is the probability that two fungi picked randomly in the community belong to different zOTUs.

Finally, to test for patterns of co-occurrence or co-exclusion of zOTUs within EPO line, we conducted three analyses. First, we tested for the correlation between zOTUs coordinates on MCA axes and *Observed index* values of each fungal clade. Then, we tested for the correlation between *Observed index* values among fungal clades. Finally, we tested for the correlation between zOTUs coordinates among fungal clades. For each analysis, we used the *cor* function of R base, and we applied a Bonferroni correction to account for multiple testing.

### Quantitative genetic analyses

#### Spatial modeling and heritability computation

To account for variation in mycobiota community composition due to variation in local environmental conditions, we fitted a spatial model with the *PSANOVA* function in the SpATS package for presence-absence of each zOTUs and each mycobiota composition variable, *i.e.* alpha diversity indexes and genotype coordinates on MCA axes (Rodríguez-Álvarez *et al*., 2018). We used a linear logistic model with logit link function for presence-absence data, whereas we used a linear model with a gaussian linked function for the mycobiota composition variables.

We estimated the heritability of presence-absence of zOTUs and mycobiota composition variables using the function *get_Heritability*. The genotype Best Linear Unbiased Predictors (BLUPs referred as to Y in the next paragraph) was extracted from the coefficients provided by the SpATS package (Rodríguez-Álvarez *et al*., 2018).

#### GWAS

##### Models

To test for the association between variation in each mycobiota composition variable and allelic variation at each SNP marker, we used the following mixed model (Yu *et al*., 2006):

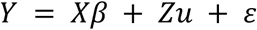

with Y the analyzed phenotype, u is the random polygenic effect, *β* represents the vector of fixed effects (intercept, dosage of the tested SNP), X and Z are the incidence matrices for fixed and random effects, respectively.

Variances of random effects were estimated as follows:

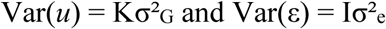

Where K represents a matrix of genetic relatedness between individuals (see below), I is the identity matrix, σ²_G_ and σ²_e_ are the polygenic and error variances, respectively. K was computed following (VanRaden, 2008) equation:

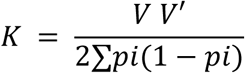

With V, the centered marker matrix and pi the allele frequency at marker i.

Each SNP was tested successively with the function *GWAS* from the R package Statgeng was based on the method presented in (Kang *et al*., 2010), using the EMMA algorithm. Indeed, this algorithm allows estimating the variance of the residual and polygenic effects only once by fitting a model with no SNP. QQ-Plots were systematically inspected to control for false positives.

##### Identification of significant SNPs

To identify significant SNPs linked to variation in mycobiota composition, we used a threshold of 4.34 corresponding to a Bonferroni type threshold computed with the Galwey method (Galwey, 2009). This threshold is well adapted to detect QTLs with large effects.

##### Computation of QTL boundaries

To group all significant markers into QTLs and thus define QTL boundaries for each mycobiota composition variable, we used a method inspired from (Cormier *et al*., 2014). For each chromosome, we computed linkage disequilibrium (LD) between significant markers, using R^²^ estimator from (Hill & Robertson, 1968). LDs were square roots transformed to approximate a normally distributed random variable as in (Breseghello & Sorrells, 2006). Then, markers were clustered by LD blocks. Clustering was performed with the UPGMA method using a cutoff of 1-“critical R²”. Critical R² (R²c) was defined as the 99.9^th^ percentile of the distribution of unlinked R² computed between 10 000 pairs of markers randomly sampled from different chromosomes. This threshold accounts for a risk of 0.1% to be in LD by chance.

QTL boundaries were defined as the minimum and maximum map positions of significant markers belonging to the same LD block. QTLs of different mycobiota composition variables were considered to overlap when they had at least one common significant marker and were located at a physical distance below one tenth of the total physical length of the chromosome as presented in (Cormier *et al*., 2014).

### Data analysis

All statistical analyses were conducted in R 4.1.

## Results

### Root mycobiota of the EPO panel

The root mycobiota of the 181 wheat inbred lines grown in the field was investigated through metabarcoding of the fungal ITS2 rDNA region. Of the 6 million reads obtained, 76.3% were retained after bioinformatic processing and clustered into a total of 533 zOTUs, with 529 identified as fungi using the UNITE database (Table S1, Fig. S1). This confirmed the specificity of primers for the fungal ITS2 region. From this set, 446 zOTUs were assigned to a specific fungal clade (*i.e.* at the phylum or subphylum rank), and retained for further analysis. The 83 remaining zOTUs were either assigned only at the kingdom level (70 zOTUs; 16.6% of the reads) or multi-affiliated (13 zOTUs; 9.1% of the read). The 446 zOTUs were distributed across 7 fungal clades, and mainly belongs to the *Ascomycota* (215 zOTUs; 38.8% of the reads) and the *Basidiomycota* (121 zOTUs; 13.8% of the reads) (Fig. 2). The remaining zOTUs belongs to the subphylum *Glomeromycotina, i.e.* AM fungi (38 zOTUs; 1.2% of the reads), the subphylum *Mortierellomycotina* (37 zOTUs; 6.1% of the reads), the phylum *Chytridiomycota* (19 zOTUs; 0.4% of the reads), the phylum *Olpidiomycota* (12 zOTUs; 13.1% of the reads), and the subphylum *Mucoromycotina* (4 zOTUs; 0.7% of the reads) (Fig. 2). The *Mucoromycotina* subphylum was excluded from further analysis due to its low number of zOTUs. Affiliations and abundance file are provided in Supplementary Information (Table S1, Table S2).

**Figure 2:**
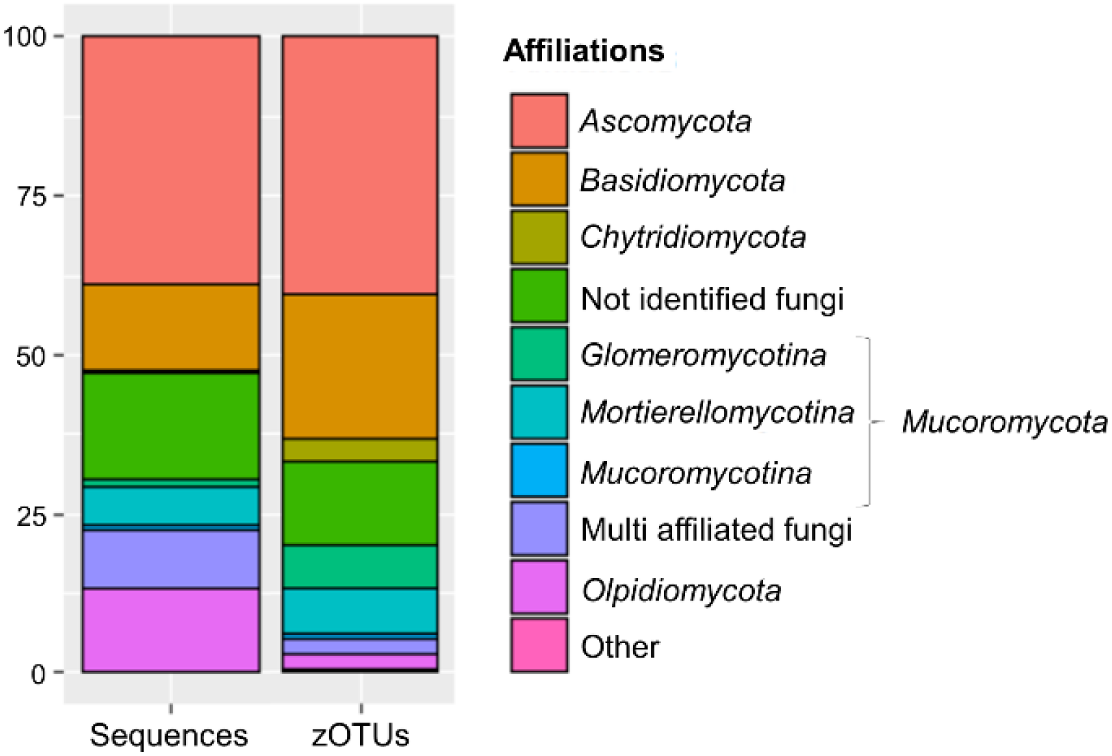
Percentage of sequences and zOTUs obtained for each fungal clade (affiliation at the phylum or sub-phylum rank) and for fungi non or multi identified. Other refers to sequences or zOTUs not affiliated as fungi.

### The root mycobiota varies depending on EPO lines

Correspondence analyses (MCA) were conducted to summarize information and assess patterns of co-occurrence or avoidance of fungal clades (*i.e.* at the phylum or subphylum rank). We analyzed the grouping of categories (presence / absence of each zOTU) along the first four axes of the MCA. For each clade, if all present zOTUs cluster together and oppose clustered absent zOTUs, it indicates that the primary determinant of fungal diversity within each sample is the presence of the entire clade rather than a specific set of zOTUs of the fungal clade.

For each of the six clades (Fig. 3) as for the whole fungal kingdom (Fig. S2), both present and absent zOTUs were grouped in the two-dimensional space along axis 1 and axis 2. Thus, for each fungal clade, the two first axes of MCA discriminate wheat root samples with either the presence or the absence of the entire clade. Such a pattern was not found for axes 3 and 4 in any fungal clade (Fig. S3).

**Figure 3:**
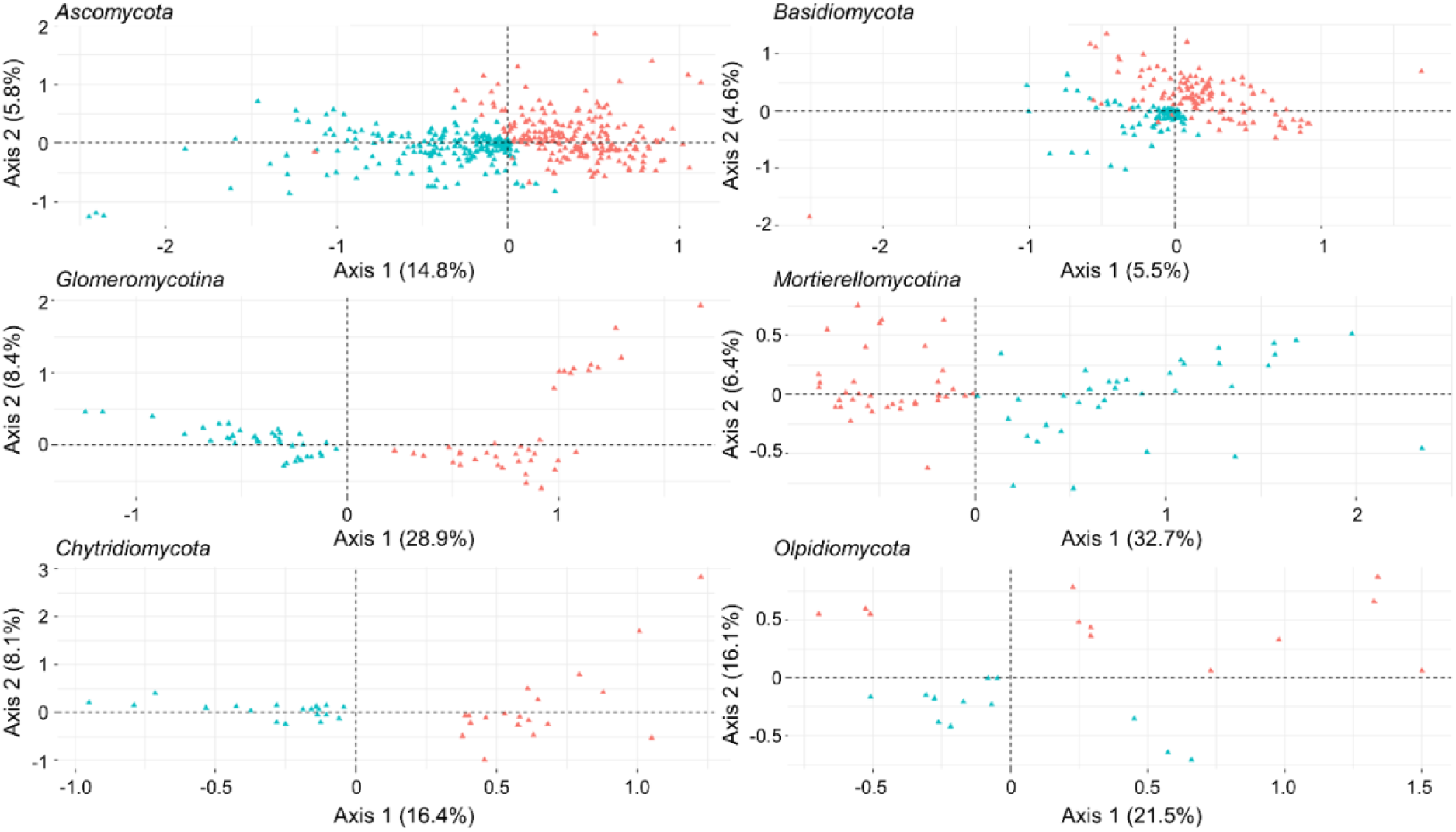
Projection of zOTUs in dimensions 1 and 2 from Multiple Correspondence Analysis for each fungal clade. Each triangle represents a zOTU, in red the projected position of its presence, and in blue the projected position of its absence.

When further testing for co-occurrence of zOTUs within EPO lines, we measured correlations between the zOTU coordinates on the MCA axes and diversity metrics for each and between clades (Fig. S4). When comparing within clades, the strongest correlations were found for axes 1 and 2 with the *Observed index* (respectively r _Glomero axe 1-Ob_ = 1; r _Asco axe 1-Ob_ = 1; r _Basidio axe 1-Ob_ = 0.8; r _Mortierello axe 1-Ob_ = −1; r _Chytridio axe 1-Ob_ = 1; r _Olpidio axe 2-Ob_ = 0.9). This confirmed that for all fungal clades, the main characteristics accounted for by the first two axes of the MCAs was either a common presence or a common absence of their respective zOTUs within each EPO line.

When comparing between clades, values of *Observed index* measured were mostly positively correlated (Fig. 4): wheat genotypes harboring high diversity in one fungal clade also exhibited high diversity in another fungal clade. Strong positive correlations were found between the *Ascomycota* and the *Basidiomycota* (r = 0.50, p<0.05) and between the *Chytridiomycota* and the *Basidiomycota* (r = 0.50, p<0.05). Interestingly, one negative correlation was detected between *Observed index* of the *Mortierellomycotina* and the *Glomeromycotina* (r = −0.30, p<0.05): genotypes with high diversity in *Mortierellomycotina* exhibited low diversity of *Glomeromycotina*, and vice versa.

**Figure 4:**
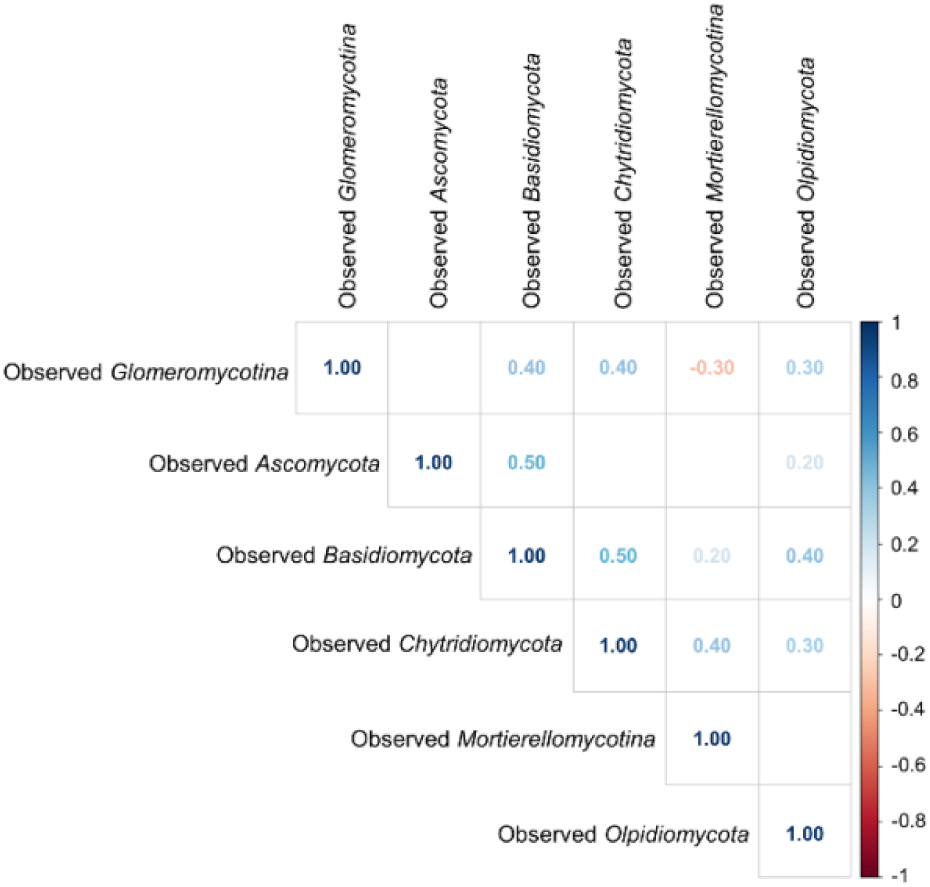
Correlation matrix of *Observed index* values for each pair of fungal clades. Only significant correlations were reported. Positive and negative correlations are represented by blue and red values, respectively. Bonferroni correction was applied to account for the 15 simultaneous testing which correspond to a critical r of 0.217. Critical r corresponds to r value above which the correlation is significant.

We found additional significant correlations between MCA axis 1 or 2 of different fungal clade (Fig. S4). As these metrics represent the presence/absence of each clade (Fig.3), such correlations could be interpreted as co-occurrence or co-exclusion between fungal clades, confirming the correlation shown with the *Observed index*. It should be noted that the sense of projection red-presence/blue-absence in Figure 3 has no biological signification (opposite sense in axis 1 of *Mortierellomycotina* clade compared to other clades). Differently from other fungal clades, the MCA axis 2 of *Olpidiomycota* and not the axis1 allows discriminating the presence-absence (Fig.3). As represented in figure S4, we found significant co-occurrence of *Olpidiomycota* and *Mortierellomycotina* (r _Olpidio axe 2 - Mortierello axe 1_ = −0.5), *Ascomycota* and *Basidiomycota* (r _Basidio axe 1 - Asco axe 1_ = 0.4), *Chytridiomycota* and *Glomeromycotina* (r _Chytridio axe 1 - Glomero axe 1_ = 0.4), *Chytridiomycota* and *Olpidiomycota* (r _Chytridio axe 1 - Olpidio axe 2_ = 0.3), *Chytridiomycota* and *Mortierellomycotina* (r _Chytridio axe 1 - Mortierello axe 1_ = −0.3). We also found a significant co-exclusion between *Mortierellomycotina* and *Glomeromycotina* (r _Mortierelle axe 1 - Glomero axe 1_ = 0.3).

### Genetic determinants of root mycobiota composition

The genetic determinants of variables describing the mycobiota composition were explored on three distinct datasets: presence-absence of zOTUs, genotype coordinates on MCA axes and the diversity indices. To account for variation in mycobiota composition due to variation in local environmental conditions, we corrected the data using a spatial model (see methods).

As a prerequisite, we assessed the heritabilities of each variable within each clade and across all clades. Heritabilities of presence-absence of individual zOTUs were weak in all fungal clades, ranging from 0.01 to 0.21 (Fig. **5a**). Among the investigated zOTUs, 113 (=25%) exhibited presence-absence features that were partially influenced by the plant genotype, as indicated by non-zero values.

**Figure 5:**
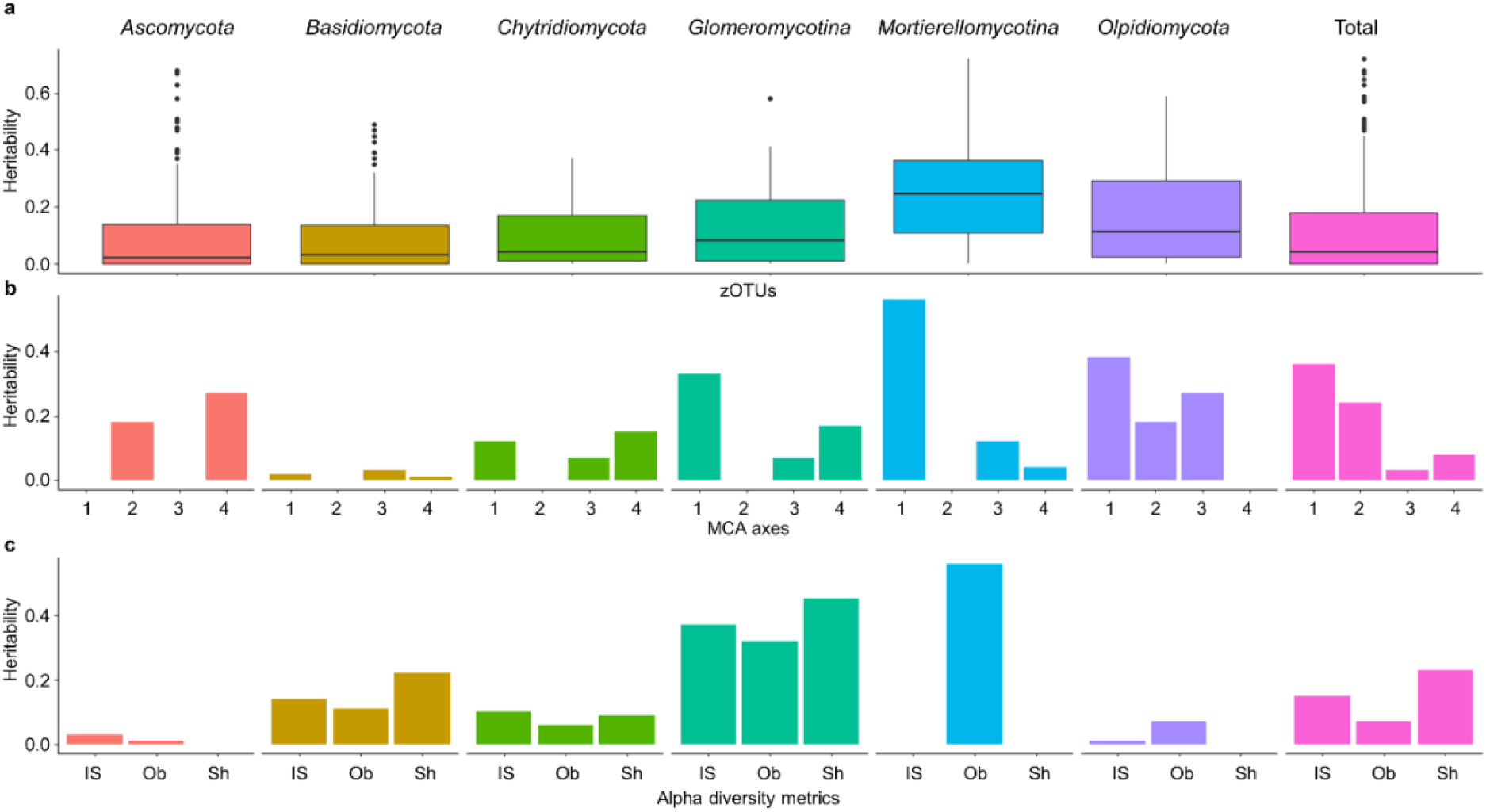
Heritabilities of mycobiota composition variables for each fungal clade and across all clades. A) Boxplot of heritabilities absence/presence of individual zOTUs. B) Heritabilities calculated with genotype coordinates obtained from the four first MCA axes. C) Heritabilities calculated with Alpha diversity metrics (*Inverse Simpson (IS), Observed (Ob) and Shannon (Sh) indexes*).

The heritabilities of genotype coordinates on MCA axes and diversity indices, were generally much higher (Fig. **5b,c**). Notably, the heritabilities of the first MCA axes for the *Mortierellomycotina,* the *Olpidiomycota* and the *Glomeromycotina* were high compared to other phyla (*h²_Mortierello_ = 0.56, h²_Olpidio_ = 0.38, h² _Glomero_= 0.33*; Fig. **5b**). Finally, the heritability of the first and second axes of the global MCA was moderate (*h²_Total axe 1_ = 0.36, h²_Total axe 2_ = 0.24*). The heritability of *Observed index* was 0.56 for the *Mortierellomycotina* and 0.32 for the *Glomeromycotina*, heritability for the *Shannon index* for the *Glomeromycotina* was 0.45 (Fig. **5c**). Overall, indices describing the *Glomeromycotina* communities had much higher values of heritabilities than other fungal clades.

To identify QTLs associated with mycobiota composition, we performed GWAS on the variables with the highest heritabilities (i.e. genotype coordinates on the MCA axes and diversity indices). We found 11 significant QTLs (Table S3). QTLs were detected in all fungal clades, although the metrics for QTL detection varied among clades. The most significant QTL was observed for the first axis of the *Chytridiomycota* (-log10(pval) = 5.4). Additionally, some QTLs were shared across multiple variables. For instance, the QTL associated with the *Shannon index* of the *Glomeromycotina* on chromosome 1A was also detected for the second axis in the global analysis.

## Discussion

In addition to recognized environmental factors playing a major role in shaping root microbiomes, there is a growing interest in exploring to which extent plant genetics also influences microbiomes (Bergelson *et al*., 2021; Raajmakers & Kiers, 2022; Escudero-Martinez & Bulgarelli, 2023). While the majority of published studies have predominantly focused on bacteria, we here studied wheat root associated fungi. Consistent with previous findings (Zheng *et al*., 2021), the wheat root mycobiota identified in our work exhibited a wide range of lifestyles, encompassing mutualists and pathogens, but also numerous fungi often described as saprotrophs. It is worth noting that our procedure to enrich the representativity of endophytic fungi in data did not fully eliminate fungi from other compartments as rhizoplane rhizosphere or soil. Although rare, some soil contaminant fungi – *i.e.* not endophytic –are present in our zOTUs, albeit in low abundance (estimated to lower than 0.5%). Overall, the robustness of our analysis is supported by the presence of the majority of fungal species already recognized as wheat endophytic fungi (Błaszczyk *et al*., 2021).

We investigated the influence of wheat genetics on root mycobiota composition by analysing both the patterns of covariation between taxa from different fungal clades, and classic quantitative genetics methodology (heritability and GWAS study). Recent studies showed an effect of domestication and modern breeding on root bacterial and fungal microbiota composition (Kinnunen-Grubb *et al*., 2020; Spor *et al*., 2020; Tkacz *et al*., 2020; Mauger *et al*., 2021; Abdullaeva *et al*., 2024). To maximize the chance to identify genetic determinants controlling microbiota composition, it is thus important to use a genetic material containing alleles potentially lost in modern cultivars. The 181 EPO lines, obtained from a wheat population with the introgression of alleles from wild durum wheat genotypes, offered this potentiality.

We provide evidence that the mycobiota composition is influenced by the genotype of the plant in wheat. Indeed, positive correlations of observed index of each detected clade suggests that the ability to find a high or low diversity of fungi in a wheat line depends on its genetics.

Using the EPO lines, we identified several QTLs associated with descriptors of mycobiota composition at the fungal phylum/sub-phylum level. However, the effect sizes were generally weak, and no QTL was identified for the majority of the descriptors, as previously reported (Zancarini *et al*., 2021). The results are in lines with studies on other species. For instance, the rhizosphere microbiome of maize (Peiffer *et al*., 2013) and sorghum (Deng *et al*., 2021) have been shown to include heritable bacterial taxa, with numerous QTLs identified, but only for a limited set of diversity descriptors. Overall reported proportions of variance explained by these QTLs are weak and heritabilities were also modest. Given the reported beneficial effects of root-associated mycobiota on crop tolerance to abiotic stress (Andreo-Jimenez et al., 2023), it would have been interesting to investigate the composition of the EPO root mycobiota under a range of abiotic field conditions. Nonetheless, by elucidating the genetic regulation of fungal diversity and identifying key fungal associations, this study advances our understanding of plant-mycobiota interactions and highlights the potential for breeding strategies aimed at optimizing these beneficial relationships (Hartman *et al*., 2018; Kavamura *et al*., 2021).

Notable differences were found between fungal clades. Taxa composition of the clades *Mortierellomycotina*, *Glomeromycotina* and *Olpidiomycota* were among those with the highest heritability. *Mortierellomycotina* and *Glomeromycotina* are phylogenetically sister clades (Spatafora *et al*., 2016), both recognized for their plant growth-promoting effects. In contrast, *Olpidiomycota*, described in the literature as soil-borne obligate parasites of Brassicaceae (Lay *et al*., 2018), were abundantly found in our root samples. *Olpidiomycota* were also previously identified in wheat (Zheng *et al*., 2021), suggesting that they might have an additional endophytic lifestyle. In contrast, composition of fungal clades containing species with a wide variety of trophic modes, such as *Ascomycota* or *Basidiomycota*, were less heritable. We analysed heritability of subgroups of *Ascomycota* or *Basidiomycota* taxa according to their supposed lifestyle (Nguyen *et al*., 2016), but it did not lead to higher heritability, probably due to the fragmentary knowledge on the lifestyle of soil fungi. Altogether heritability results suggest that the influence of host genetic control on root mycobiota taxa should partly rely on the dependency of endophyte fungi for their host. Obligate biotrophs as *Glomeromycotina*, fully depending of host carbon resources, are much more under host genetic control than facultative endophytes that have to share nutrient resources from root exudates or from apoplastic fluid. Mirroring this fungal dependency to the host, since phylogenetically divergent microbial taxa are known to provide similar services, it is expected that host-genetics regulation is on services, functions and lifestyle rather than on taxa identities (Lemanceau *et al*., 2017). Some general services are shared by various fungi, as phytohormone synthesis, while others are more specific, as the transfer of nutrients to the host (as nitrogen or phosphorus). Thus, the genetic control by the plant of the services provided by its mycobiota could be higher on fungal clades having a specific life style and providing a limited number of services, such as the mutualistic *Glomeromycotina*, than on large clades such as the *Ascomycota* and the *Basidiomycota* which supply a large number of diversified and redundant services. Another possibility is that the services provided by *Mortierellomycotina*, *Glomeromycotina* and *Olpidiomycota*, under the experimental conditions applied here, were more important. A future challenge would thus be to identify host genetic determinants controlling mycobiota services rather than mycobiota composition. This would first require to functionally characterize the fungal diversity, which is far to be the case in spite of existing tools such as FUNGuild (Nguyen *et al*., 2016) and FungalTraits (Põlme *et al*., 2020), due to the cryptic lifestyle of most fungi. Finally, beside understanding host control over microbiota composition/functions, it will be essential to study the degree of host genetic variability to benefit from the root microbiota composition/functions, in order to exploit this variability in breeding programs.

Interestingly, complex relation exists between fungal clades associated with wheat root. A negative correlation was found between zOTU richness from the *Mortierellomycotina* and from the *Glomeromycotina*, suggesting some compensatory effect. The AM fungi *Glomeromycotina* are ubiquitous mutualistic fungi associated with most of the plant roots (Smith & Read, 2008). *Mortierella* sp., belonging to *Mortierellomycotina,* were described as phosphate solubilizing microorganisms (Zhang *et al*., 2011). Both are known to promote plant growth especially by enhancing plant nutrition (Zhang *et al*., 2020). Inoculation of these fungi individually has shown a positive impact on wheat growth and yield in both field and pot experiments (Singh & Reddy, 2011; Berruti *et al*., 2015; Ozimek *et al*., 2018). Inoculation of both fungi has been less documented especially in agricultural soil. Velez & Osorio (Tamayo-Velez & Osorio, 2017) have tested co-inoculation of *Mortierella sp.* and AM fungal isolates on avocado plantlets in controlled conditions. Interestingly, they found an additive effect of dual inoculation on shoot avocado height and P content and a decrease of both fungal root colonization suggesting that AM fungi and *Mortierellomycotina* species might share the same ecological niche. The most straightforward hypothesis posits that both clades confer similar benefits to their host such as phosphorus uptake. The observed exclusion pattern in our analysis may consequently arise from strong competition, driven by a process of species exclusion vying for the same endophytic niche.

Besides this compensatory/negative effect, positive correlations have been found between zOTU richness for other pairs of fungal clades, among which the *Chytridiomycota*/*Basidiomycota* is the highest. This could suggest synergistic or facilitative interactions among particular species within these fungal clades. Alternatively, these positive correlations could demonstrate a general capacity of certain plant genotypes to accommodate fungi within their tissues. This implies potential mechanisms associated with quantitative control of fungal invasion, whether correlated with or independent of plant immunity. Describing precisely these interactions would require sophisticated statistical analyses and integration of multiple data types (Zancarini *et al*., 2021).

In conclusion, our study demonstrates that wheat genotypes significantly influence the composition of the root mycobiota. The apparent ability of specific genotypes to host a limited fungal diversity merits further investigation, paving the way for the identification of novel quantitative plant resistance mechanisms. Regarding the mycobiota, our observations emphasize the importance of evaluating the role of *Olipidiomycota* and *Chytridiomycota* as root endophytes. Furthermore, our findings emphasize the potential of AM fungi and *Mortierellomycotina* as promising fungal associations with potential benefits as plant-growth promoting fungi.

## Acknowledgements

We thank Antonin Grau (EU DIASCOPE, INRAE) for supervision of the EPO field assay. We also thank Virginie Gasciolli and Coline Rosset for their technical assistance. We are grateful to Jean-François Allienne, Margot Doberva and Michèle Laudié from the Bio-Environment platform (UPVD, Région Occitanie, CPER 2007-2013 Technoviv, CPER 2015-2020 Technoviv2) for technical support in library preparation and sequencing.

## Authors’ contributions

CR, BL, JD & HF designed the experiment. AR set up the field assay. MT produced and analysed the Miseq data. MC performed GWAS analysis under JD supervision. MT & MC wrote the manuscript under the supervision of CR, BL, JD & HF.

JD, HF and MC were supported by the Agence Nationale de la Recherche (ANR) project “Selecting for cooperative crops to develop sustainable agriculture” (SCOOP, grant no.ANR-19-CE32-0011). CM is also co-funded by INRAE-Département “Biologie et Amélioration des Plantes”. This study is set within the framework of the “Laboratoires d’Excellences (LABEX)” TULIP (ANR 10 LABX 41) and of the “Ecole Universitaire de Recherche (EUR)” TULIP GS (ANR 18 EURE 0019). MT PhD is co-funded by the Université Fédérale Toulouse Midi-Pyrénées (UFTMiP) and the Région Occitanie.

## Data availability

All data will be available under request.

