## Supplementary figures 1-4 for "Host genotype shapes root mycobiota in durum wheat"

| | Total sequences number | Total zOTU number |
|---|---|---|
| **Imput sequences** | 6 086 748 | - |
| **Pre-process** | 5 348 156 (87,9%) | - |
| **Clustering** | 5 348 156 | 208 352 |
| **Remove chimera** | 5 211 004 (97,4%) | 165 969 (79,7%) |
| **zOTU Filter** | 4 898 536 (94%) | 552 (0,3%) |
| **ITS Filter** | 4 652 599 (95%) | 533 (96,6%) |
| **Affiliated zOTU** (at kingdom level) | 4 645 690 (99,9%) | 529 (99,2%) |
| **Affiliated zOTU** (at species level) | 2 679 606 (57,7%) | 242 (45,7%) |

Supp. Figure 1: Number of sequences and zOTUs retained throughout the bioinformatics processing. In blue: the proportion of sequences or zOTUs that persist compared to the preceding step.

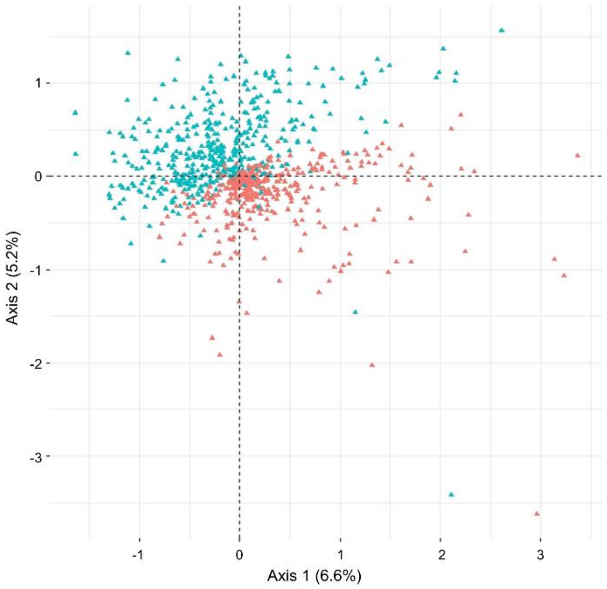


Supp. Figure 2: Projection of zOTUs in dimensions 1 and 2 from Multiple Correspondence Analysis of the global dataset. Each triangle represents a zOTU, in red the projected position of its presence, and in blue the projected position of its absence.

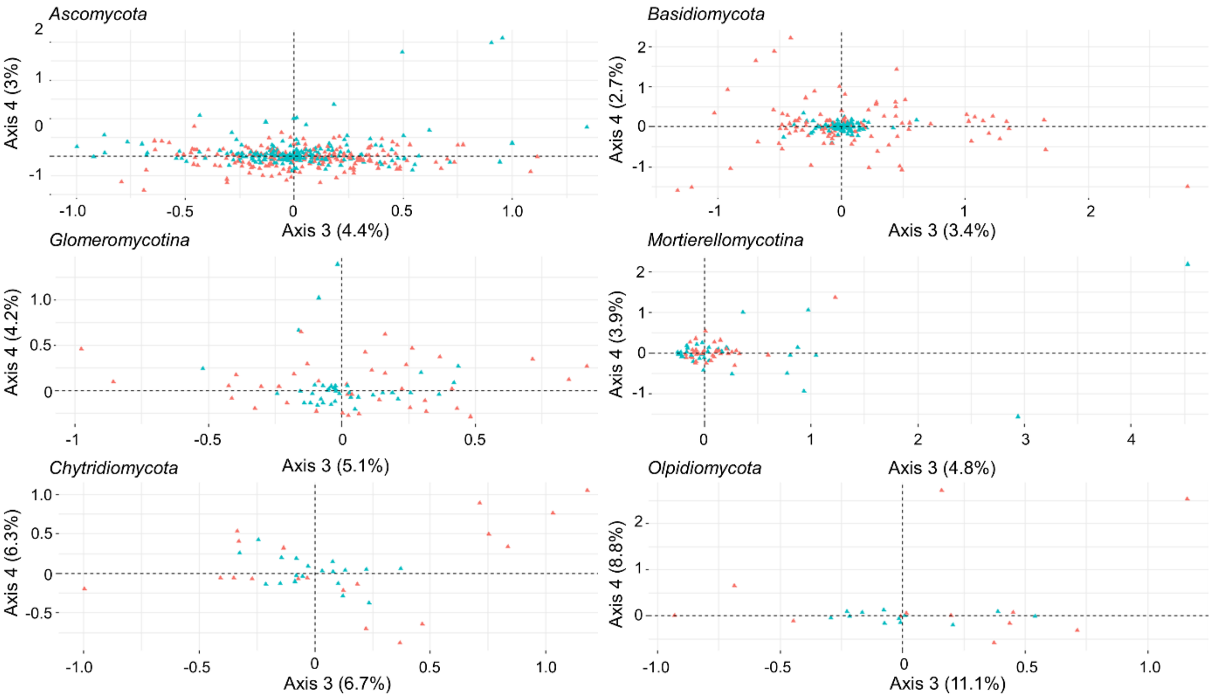


Supp. Figure 3: Projection of zOTUs in dimensions 3 and 4 from Multiple Correspondence Analysis for each fungal clade. Each triangle represents a zOTU, in red the projected position of its presence, and in blue the projected position of its absence.

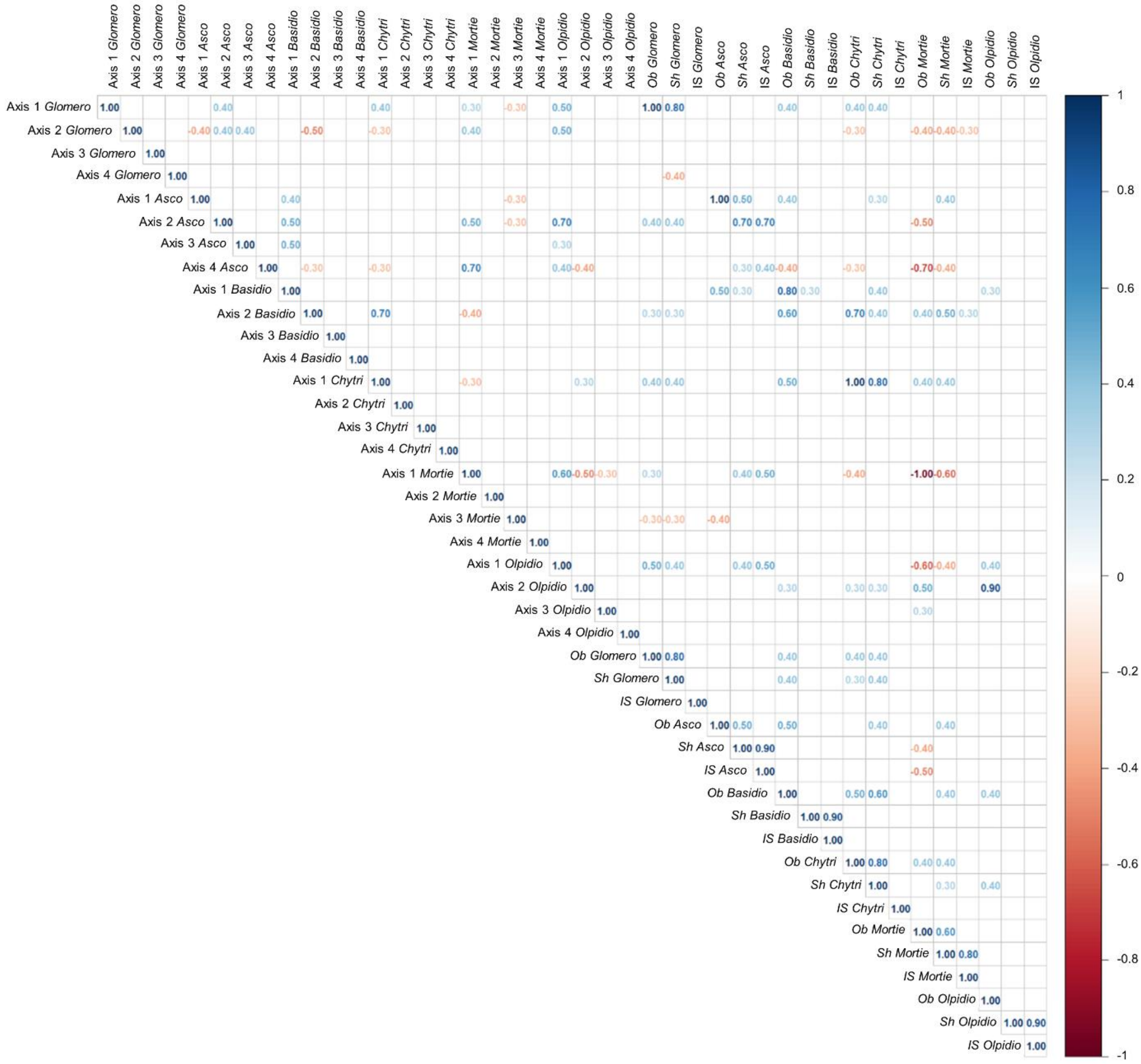


Supp. Figure 4: Correlation matrix between all mycobiota composition variables for each pair of fungal clades. Only significant correlations were reported. Positive and negative correlations are represented by blue and red values, respectively. Bonferroni correction was applied to account for the 861 simultaneous testing which correspond to a critical r of 0.3.
